# A viral vaccine design harnessing prior BCG immunization confers protection against Ebola virus

**DOI:** 10.1101/2024.05.28.595735

**Authors:** Tony W. Ng, Wakako Furuyama, Ariel S. Wirchnianski, Noemí A. Saavedra-Ávila, Christopher T. Johndrow, Kartik Chandran, William R. Jacobs, Andrea Marzi, Steven A. Porcelli

## Abstract

Previous studies have demonstrated the efficacy and feasibility of an anti-viral vaccine strategy that takes advantage of pre-existing CD4^+^ helper T (Th) cells induced by *Mycobacterium bovis* bacille Calmette-Guérin (BCG) vaccination. This strategy uses immunization with recombinant fusion proteins comprised of a cell surface expressed viral antigen, such as a viral envelope glycoprotein, engineered to contain well-defined BCG Th cell epitopes, thus rapidly recruiting Th cells induced by prior BCG vaccination to provide intrastructural help to virus-specific B cells. In the current study, we show that Th cells induced by BCG were localized predominantly outside of germinal centers and promoted antibody class switching to isotypes characterized by strong Fc receptor interactions and effector functions. Furthermore, BCG vaccination also upregulated FcγR expression to potentially maximize antibody-dependent effector activities. Using a mouse model of Ebola virus (EBOV) infection, this vaccine strategy provided sustained antibody levels with strong IgG2c bias and protection against lethal challenge. This general approach can be easily adapted to other viruses, and may be a rapid and effective method of immunization against emerging pandemics in populations that routinely receive BCG vaccination.

## 1 Introduction

Emergent viruses such as Ebola virus (EBOV) and related filoviruses are global health threats of increasing concern, especially due to the expansion of human populations into wild habitats that serve as natural reservoirs for these viruses ^1^. For prevention of outbreaks of viral infections or pandemics, vaccines remain the most practical and cost-effective tools. This has clearly been shown in the ongoing Coronavirus disease 2019 (COVID-19) pandemic where vaccination has been reported to reduce the risk of severe illness leading to hospitalization and mortality rates among vaccinated individuals ^2^. Historically, vaccine development has been mainly focused on the variable (Fab) region of immunoglobulins for their ability to bind surface antigens of viruses and prevent entry into host cells ^3–5^. Such neutralizing antibodies (NAbs) are a major correlate of protection associated with viral clearance and resolution of the infection. However, the limited range of epitopes available to induce NAbs may prevent efficient clearance of infection by this mechanism for some viruses. Furthermore, as shown in human immunodeficiency virus (HIV) infection, NAbs exert strong selective pressure in driving immune escape of the virus as compared to non-neutralizing antibodies ^6^. These observations suggest that eliciting non-neutralizing antibodies mediating effector functions distinct from simple blockade of viral entry might increase the protective efficacy of a vaccine ^7, 8^.

The constant (Fc) regions of antibodies, although recognized as important in contributing to protection, have been less emphasized compared to the Fab region as a determinant of anti-viral effects. The Fc region binds to the Fc receptors (FcRs) on a variety of relevant immune effector cells, thus bridging humoral and cellular immunity through effector activities such as antibody-dependent cellular phagocytosis (ADCP), complement-dependent cytotoxicity (CDC), and antibody-dependent cellular cytotoxicity (ADCC) that are believed to contribute to control of viral and other microbial infections ^9, 10^. Recently, results from clinical trials for newly developed EBOV and HIV vaccines have called attention to the importance of antibody-mediated effector functions as correlates of protection against viral pathogens ^11–15^. This has driven the search for relevant antibody effector functions beyond simple neutralization in individuals vaccinated against or exposed to EBOV ^16, 17^ or HIV ^18, 19^, and has encouraged efforts to engineer therapeutic antibodies with optimal Fc effector functions for these diseases ^20, 21^.

Whereas the binding affinity of the Fab region of the antibody develops and matures in the germinal centers (GC) within B cell follicles of secondary lymphoid tissues ^22^, class-switch recombination (CSR) required for determining Fc isotype is initiated and occurs mostly at the border of the B cell follicle between the boundary of B and T cell zones ^23^. Activation of CSR requires signals from B cell receptor (BCR) engagement, costimulatory signals such as CD40-CD40L interaction, and particularly cytokines secreted from CD4^+^ helper T cells (Th) that dictate which switch region of the heavy chain constant region genes will interact with activation-induced cytidine deaminase (AID) to initiate the double strand DNA break required for recombination to occur ^24^. Therefore, the ability to induce different Th phenotypes, such as Th1, Th2 or follicular helper T cells (Tfh), during vaccination can have an impact on class-switching of immunoglobulins. In C57BL/6 mice, for example, class-switching to IgG2c (homologous to IgG2a in other mouse strains ^25^), is induced by IFNγ produced by Th1 cells ^26, 27^, whereas IgG1 is induced by IL-4 derived mainly from Th2 cells ^28–30^. These antibody subclasses have different affinities to particular Fc receptors (FcRs) ^31^. In mice, antibodies with the IgG1 isotype have low, but similar affinities for the inhibitory FcγRIIB and the activating FcγRIII, whereas the affinities conferred by the IgG2 isotypes for the activating FcγRIV are much stronger ^31^. Under inflammatory Th1 conditions, IgG2c class switching and an increased expression of FcγRIV are favored ^32^, thus promoting effector functions such as phagocytosis, complement activation and cytotoxicity, all contributing to the removal of either the pathogen itself or the cells infected by it.

*Mycobacterium bovis* bacille Calmette-Guérin (BCG), the only currently approved vaccine against tuberculosis, is one of the most widely administered vaccines in many regions of the world. The BCG vaccine induces long lasting BCG-specific memory CD4^+^ T helper cells (Th) that are strongly polarized to IFNγ-secreting Th1 cells in vaccinated individuals. To take advantage of the high prevalence of BCG vaccination, we developed a vaccination strategy that uses pre-existing BCG-specific Th cells to drive antibody responses against a modified viral protein immunogen ^33^. This recombinant fusion protein vaccine (Th-vaccine), based on a principle previously designated intrastructural help ^34, 35^, induces antiviral antibody responses with a strong bias to IgG2c isotype in C57BL/6 mice ^33^. In the current study, we have used an EBOV challenge model to show that this approach promotes antibodies that can recruit effector cells capable of eliminating virus infected cells and is highly effective at protecting mice from lethal virus challenge. Analysis of the underlying mechanism for the effects on the antibody response showed that BCG vaccination created inflammatory conditions that impaired GC formation, similar to what has been seen in infections with other Th1 skewing pathogens ^36–39^. Antibody class switching thus occurred outside of the GC, leading to a strong bias of anti-EBOV antibodies to IgG2c isotype due to the influence of BCG-specific Th1 cells. The anti-EBOV GP antibodies elicited by this vaccination regimen were maintained over time suggesting the induction of long-term protection. Overall, our findings support the importance of non-neutralizing antibodies in anti-viral vaccination, and define a powerful and potentially useful method to induce such antibodies against established or newly emerging viruses in populations that receive routine BCG vaccinations.

## 2 Materials and Methods

### Mice

Five-week old female wild-type (WT) C57BL/6NHsd and C57BL/6J mice were obtained from Envigo (Greenfield, IN) and The Jackson Laboratory (Bar Harbor, ME), respectively. The GFP^+^ C57BL/6-P25 TCR transgenic (Tg) mice ^33^ with T cell receptor that recognizes the P25 peptide (FQDAYNAGGHNAVF) from *M. tuberculosis* or BCG Ag85B were maintained and bred in our facility. All mice were maintained in our specific pathogen-free facilities following protocols and regulations established by the Albert Einstein College of Medicine Institutional Animal Use and Care and the Institutional Biosafety Committees. All procedures performed on these animals were approved by the Albert Einstein College of Medicine Institutional Animal Use and Care Committee.

### Mycobacterial strains and vaccinations

*Mycobacterium bovis* BCG Danish strain (Statens Serum Institut, Copenhagen, Denmark) was the BCG vaccine strain used in this study. Starting from a low-passage-number frozen stock, the bacteria was grown at 37°C shaking in Sauton medium until mid-log phase, centrifuged at 600 x *g* for 10 minutes, and resuspended in sterile PBS (Thermo Fisher Scientific, Waltham, MA). BCG was administered by subcutaneous (s.c.) injection at the scruff of the neck at a dose of 1 x 10^7^ CFU. For recombinant protein vaccines injections, the vaccine in PBS was mixed in a 1:1 volume ratio with alum suspension (Imject Alum; Thermo Fisher Scientific) to a final concentration of 0.5 µg/ml unless otherwise specified, and 100 µl was administered intramuscular (i.m.) into the thigh muscles with 50 µl per hind limb to provide the final dose of 0.05 µg of the recombinant protein vaccine per animal.

### Cell lines

FreeStyle 293-F cells (Thermo Fisher Scientific) were maintained in Life Technologies FreeStyle 293 Expression Medium with GlutaMAX (Thermo Fisher Scientific). Murine T cell hybridomas (TCHs) specific for I-A^b^ -restricted CD4 T cell epitopes ^33^ were maintained in complete RPMI (cRPMI) which consists of RPMI 1640 (Thermo Fisher Scientifics) supplemented with 10 mM HEPES, 50 µg/ml penicillin/streptomycin, 55 µM 2-mercaptoethanol (Thermo Fisher Scientifics), and 10% heat-inactivated [56°C, 30 min] fetal bovine serum (Atlanta Biologicals, Flowery Branch, GA).

### Plasmid construction

The recombinant protein vaccine that consists of the extracellular portions of EBOV GP lacking the MLD (WT GPΔM) and a similar version that consists of BCG Th epitopes fused to the N terminus of the EBOV GP (Th GPΔM) were constructed and described previously ^33^. The full-length EBOV GP with the MLD restored was also constructed for these recombinant protein vaccines. To construct the full-length version of the EBOV GP vaccines, the MLD of the EBOV GP was amplified from pMAM01 ^40^ using primer pair TN258 (5΄- GCGCACCGTCGTGTCAAACGGAGCCAAAAACATCAGTGG-3΄) and TN259 (5΄- GCGCCAGTATCCTGGTGGTGAGTGTTGTTGTTGCCAGCGG-3΄). The MLD was cloned into WT GP-ΔM and Th GP ΔM via the AleI and XcmI sites to create the corresponding version WT GP-FL and Th GP-FL which contain the MLD in the EBOV GP.

### Expression and purification of recombinant protein vaccines

Plasmids corresponding to WT GP-FL and Th GP-FL DNA were transfected into FreeStyle 293-F cells, and proteins were collected from culture supernatants and purified using the HisTrap HP column (GE Healthcare Life Sciences, Pittsburgh, PA) as described previously ^33^. Protein concentrations were determined by the bicinchoninic acid (BCA) assay (Thermo Fisher Scientific). The purified proteins (220ug/ml WT GP-GL and 80ug/ml Th GP-FL) in PBS were stored at −80°C until needed. Purified Th vaccines were analyzed by size-exclusion high performance liquid chromatography (SE-HPLC) using the SRT SEC-300 size exclusion column. Analysis of the molecular weight was determined by comparing the retention time with markers of known molecular weight (BioRad).

### SDS-PAGE analysis of recombinant protein vaccines

Purified recombinant protein vaccines were analyzed on SDS-PAGE by staining with GelCode Blue Safe Protein Stain (Thermo Fisher Scientific). Proteins separated by SDS-PAGE were also transferred onto nitrocellulose membranes for immunoblotting. After blocking with 5% bovine milk in PBS with 0.05% Tween 20 (PBST), the nitrocellulose membranes containing the purified recombinant fusion proteins were incubated with mouse anti-His antibody [clone HIS.H8] (Millipore Sigma, Burlington, Massachusetts). HRP-conjugated rabbit anti-mouse IgG antibody (SouthernBiotech, Birmingham, AL) were used as detection Abs, and signals were detected using the SuperSignal West Pico PLUS Chemiluminescent Substrate (Thermo Fisher Scientific).

### T cell hybridoma stimulation assays

Mouse T cell hybridomas specific for peptide P25 of Ag85 or peptide P10 of TB9.8 were cocultured with murine bone marrow-derived dendritic cells ^33^ and incubated with the purified recombinant protein vaccine (10 µg/ml) at 37°C for 18 h. Cell culture supernatants were assayed for IL-2 using capture and biotin-labeled detection antibody pairs (BD Biosciences, Franklin Lakes, NJ). Detection was performed with HRP-conjugated streptavidin (BD Biosciences) followed by the addition of the Turbo 3,3’,5,5’-tetramethylbenzidine (TMB) substrate (Thermo Fisher Scientific).

### ELISA assays for anti-EBOV GP antibody titers

Measurement of anti-EBOV GP antibody titers was performed by direct solid phase ELISA. Corning 96-well flat bottom assay plate (Thermo Fisher Scientific) were coated with ∼95,000 infectious unit (IU) of recombinant vesicular stomatitis virus (rVSV) expressing EBOV GP (rVSV-EBOV) in PBS (pH7.4) overnight at 4°C ^41, 42^. The EBOV GP coated ELISA plate was washed three times with PBS and blocked with 2% bovine serum albumin (BSA) in PBS for 1 hour at RT. Serum samples from immunized mice were obtained from blood collected by retro-orbital bleed in vaccinated mice. The serum samples were diluted 1:50 in PBS for single dilution measurement or 1:20 followed by serial 1:3 dilutions for endpoint titers, and then incubated in the EBOV GP coated ELISA plate wells for 2 h at RT. The ELISA plates were then washed four times with PBS and incubated with HRP-conjugated mouse IgG1- or IgG2c-specific Abs (SouthernBiotech) for 1 h at RT. After washing four times in PBS, the signal was detected with SIGMAFAST OPD substrate (Sigma-Aldrich, St. Louis, MO) and the reaction was stopped with the addition of 0.5 M H_2_SO_4_. The absorbances for both the capture and direct ELSIA assays were measured with the Wallac 1420 VICTOR2 microplate reader (Perkin Elmer, Waltham, MA).

### Flow cytometry analysis of Th subsets and FcγR expression

Naïve CD4^+^ T cells from P25 TCR-Tg/GFP mice were purified by negative selection using a commercially available kit and following the manufacturer’s instruction (Miltenyi Biotec, Auburn, CA). During the CD4^+^ T cell purification, anti-CD44 conjugated to biotin [clone IM7] (Thermo Fisher Scientific) was added in the purification step to remove memory T cells that were present in these animals. 4 x 10^4^ purified CD4^+^ T cells in 100 µl of PBS were injected intravenously via the tail vein into WT C57BL/6 mice. Sixteen hours after injection of the CD4^+^ T cells, mice were vaccinated with 100 µl of either 1 x 10^7^ CFU of BCG in PBS or with 10 µg P25 peptide in PBS formulated with one of the following adjuvants: 1:1 volume ratio of alum (Imject Alum; Thermo Fisher Scientific), or 5% final volume of LASTS-C [Span85-Tween 80-squalene, lipid A, CpG oligodeoxynucleotides] ^43, 44^ (gift from Dr. Michael Anthony Moody, Duke University). On day 7 after vaccination, mice were sacrificed, and spleens were harvested and cells were stained with Live Dead viability dye (LD Fixable Blue; Thermo Fisher Scientific L34961) and antibodies against MHC class II (Alexa Fluor 700; BD 570802), CD4 (APC-Cy7, BD 561830), T-bet (PE-Cy7; Biolegend 644823), CXCR-5 (PE; BD 551959), and Bcl-6 (APC; Biolegend 358505), and analyzed by FACS using the 5 laser BD Biosciences LSRII Flow Cytometer, and 5 x 10^5^ events per sample were collected and analyzed using FlowJo software (BD biosciences).

For analysis of FcγR expression, splenocytes from FcγRII,III,IV α-chains knockout mice ^45^ or WT B6 vaccinated mice were processed at indicated timepoints and stained with mAbs against B220 (BUV661; BD 612972, clone: RA3-6B2), NK1.1 (BV605; BD 563220, clone: PK136), CD11c (Alexa Fluor 700; BD 560583, clone: HL3), CD11b (PE-CF594; BD562287, clone: M1/70), Ly-6G/Ly-6C (APC; eBioscience 17-5931-81, clone: RB6-8C5), Ly6-C (PerCP; Biolegend 128028. clone: HK1.4), FcγRIV (PE; BD 565615, clone: 9E9), FcγRII/III (FITC; BD 561726, clone: 2.4G2), and analyzed by FACS using the Cytek Aurora configured with five lasers, three scattering channels, and sixty-four fluorescence channels, and 1 x 10^6^ events per sample were collected and analyzed using the FlowJo software (BD biosciences).

### Analysis of germinal centers

To detect the presence of antigen specific Th cells in the secondary lymphoid tissues, spleens from vaccinated mice were sectioned and stained as described previously ^33^. Briefly, naïve CD4^+^ T cells from P25 TCR-Tg/GFP mice were transferred into mice which were then vaccinated with either 1 x 10^7^ BCG or with 10 µg P25 peptide formulated in LASTS-C as described in the analysis of Th subsets above. On day 7 after vaccination, mice were sacrificed and spleens were fixed in 10% neutral buffered formalin, paraffin embedded and sectioned. Tissue sections were stained with anti-GFP Ab (A11122; Thermo Fisher Scientific) for the presence of CD4^+^ T cells transferred from the P25 TCR-Tg/GFP mouse and counterstained with hematoxylin.

### Animal ethics statement

All infectious work with MA-EBOV was performed in the maximum containment laboratories at the Rocky Mountain Laboratories (RML), Division of Intramural Research, National Institute of Allergy and Infectious Diseases, National Institutes of Health. RML is an institution accredited by the Association for Assessment and Accreditation of Laboratory Animal Care International (AAALAC). All procedures followed standard operating procedures (SOPs) approved by the RML Institutional Biosafety Committee (IBC). Mouse work was performed in strict accordance with the recommendations described in the Guide for the Care and Use of Laboratory Animals of the National Institute of Health, the Office of Animal Welfare and the Animal Welfare Act, United States Department of Agriculture. The study was approved by the RML Animal Care and Use Committee (ACUC). Procedures were conducted in mice anesthetized by trained personnel under the supervision of veterinary staff. All efforts were made to ameliorate animal welfare and minimize animal suffering; food and water were available *ad libitum*.

### EBOV challenge in vaccinated mice

Wild-type female C57BL/6NHsd (approximately 10 weeks of age) were given 1 x 10^7^ BCG in PBS through s.c. injection at the scruff of the neck. Five weeks after exposure to BCG, the mice were primed with 0.05 µg of the purified recombinant protein vaccine (WT GP-FL or Th GP-FL) adjuvanted with alum in PBS through i.m. injection as described above. Vaccinated mice were rested for 4 weeks, followed by a homologous boost of the recombinant protein vaccine administered through the same route. Two weeks after each interval of administering the purified recombinant protein vaccine, blood was collected through retro-orbital bleed to obtain serum samples for antibody titer measurements. Four weeks after the boost, mice were shipped to Rocky Mountain Laboratories in Hamilton, MT and rested for 1 week prior to MA-EBOV challenge. Mice were infected by intraperitoneal (i.p.) injection of a lethal dose for naïve mice of 10 focus-forming units (FFU) of MA-EBOV ^46^. Five mice from each vaccinated group were euthanized on day 5 after challenge to harvest organs to determine viremia and to collect blood samples to freeze down serum samples for future analysis of anti-EBOV GP antibody responses. The remaining 10 mice from each vaccinated group were kept under observation for survival and weight loss and all surviving mice were euthanized on day 28 after challenge to collect and freeze serum samples.

### ELISPOT to detect antigen specific T and B cells

For the T cell analysis, five-week-old female WT C57BL/6J mice (n = 10) were vaccinated by s.c. injection of either PBS or BCG and rested for 5 weeks. Each group was further subdivided into 2 groups (n = 5) which received either PBS or the Th GP-FL vaccine at 0.05 μg per mouse in alum. Splenocytes were obtained two weeks later for ELISPOT assay ^47^. Briefly, the 96-well ELISPOT plate (Millipore) was prepared by coating the well with 50 μl of 10 μg/ml of anti-mouse IFNγ monoclonal capture antibody (BD Biosciences; cat. no. 551309) in PBS and allowed to incubate at 4°C for 16 hours. The ELISPOT wells were washed five times with PBST and blocked with 200 μg of cRPMI for 2 hours at room temperature. Splenocytes at 5 x 10^5^ cells per well were added along with 5 μg/ml P25 peptide or 10 μg/ml of *M. tuberculosis* (strain H37Rv) lysate for antigen stimulation and incubated at 37°C in 5% CO_2_ for 16 hours. The ELISPOT plate was washed five times with PBST and 50 μl of 1 μg/ml of the anti-mouse IFNγ monoclonal detection antibody conjugated to biotin (BD Biosciences; cat. no. 551506) in PBS was added and allowed to incubate at room temperature for 2 hours. The wells were then washed five times with PBS + 0.1% Tween-20 (PBST) and streptavidin-alkaline phosphatase (Thermo Fisher Scientific) at 1:1000 dilution in PBS was added incubated at 37°C in 5% CO_2_ for 1 hour. After a final 5 washes with PBST, the spots were developed by adding the BCIP/NBT substrate (Sigma Aldrich). The reaction was stopped by washing the wells with water and the spots were counted using an automated ELISPOT reader (Autoimmun Diagnostika GmbH, Strasbourg, Germany).

For ELISPOT quantitation of antibody secreting cells ^48^, five-week-old female WT C57BL/6 mice were vaccinated with PBS or BCG by s.c. administration of 1 x 10^7^ CFU per mouse. Five weeks after vaccination, mice were injected i.m. with 5 μg of the Th GP-FL vaccine adjuvanted with alum in PBS. Thirty-nine weeks later, the mice were sacrificed to obtain the splenocytes and bone marrow cells, which were immediately assayed by ELISPOT to detect antibody secreting B cells. The 96-well ELISPOT plate (Millipore) was prepared by coating the wells with 50 μl of 10 μg/ml of rVSV expressing EBOV GP and incubating at 4°C for 16 hours. The ELISPOT wells were washed five times with PBS and blocked with 200 μl of cRPMI for 2 hours at room temperature. Splenocytes or bone marrow cells at 10^6^ cells per well were added to the ELISPOT plate and incubated at 37°C in 5% CO_2_ for 5 hours. The ELISPOT plate was washed five times with PBS and anti-mouse IgG1 or anti-mouse IgG2c antibodies conjugated with alkaline phosphatase (Southern Biotech) at 1:1000 in PBS were added and incubated at room temperature for 2 hours. The spots were developed by adding the BCIP/NBT substrate (Sigma Aldrich). The reaction was stopped by washing the wells with water and the spots were counted using an automated ELISPOT reader (Autoimmun Diagnostika GmbH, Strasbourg, Germany).

### Comparative immunogenicity of EBOV GP vaccine constructs

Subunit vaccines against EBOV have shown potential for inducing protection against infection in several animal models and may have important advantages over virally vectored vaccines ^49^. However, optimal design of subunit vaccines regarding efficacy, potency, stability and formulation issues requires further investigation and testing ^50^. The design of the EBOV glycoprotein (GP) Th-vaccine for the current study was based on our previous work showing the general impact of incorporating immunodominant Th epitopes of BCG into a soluble version of EBOV GP from which the mucin-like domain (MLD; EBOV GPΔMLD) was deleted to direct responses against conserved epitopes important for neutralizing antibodies ^33, 40^. Here we developed a new version of the EBOV GP Th-vaccine that consisted of the full-length complete extracellular portion of the EBOV GP for direct comparison with the previous version of EBOV GPΔMLD. Although the EBOV GPΔMLD was produced with higher yields as a recombinant protein, the full-length version of the EBOV GP Th-vaccine has the advantages of more closely resembling the native protein on the viral envelope or surface of infected cells, and also could provide a greater range of epitopes for antibody targeting of membrane expressed GP. As shown schematically (Fig. 1A), the immunodominant CD4^+^ T cell epitopes of mycobacterial antigens Ag85B (P25 epitope) and TB9.8 (P10 epitope) were fused to the N terminus of the full-length EBOV GP or to a version of EBOV GPΔMLD. These were designated Th GP-FL or Th GP-ΔM, respectively. Protein expression was done in FreeStyle 293-F cells and purified by Ni-NTA affinity chromatography as described previously ^33^. Versions of these proteins lacking the N-terminal extension encoding the BCG Th epitopes, designated as wild type (WT), were also constructed and purified to serve as controls.

**Figure 1.**
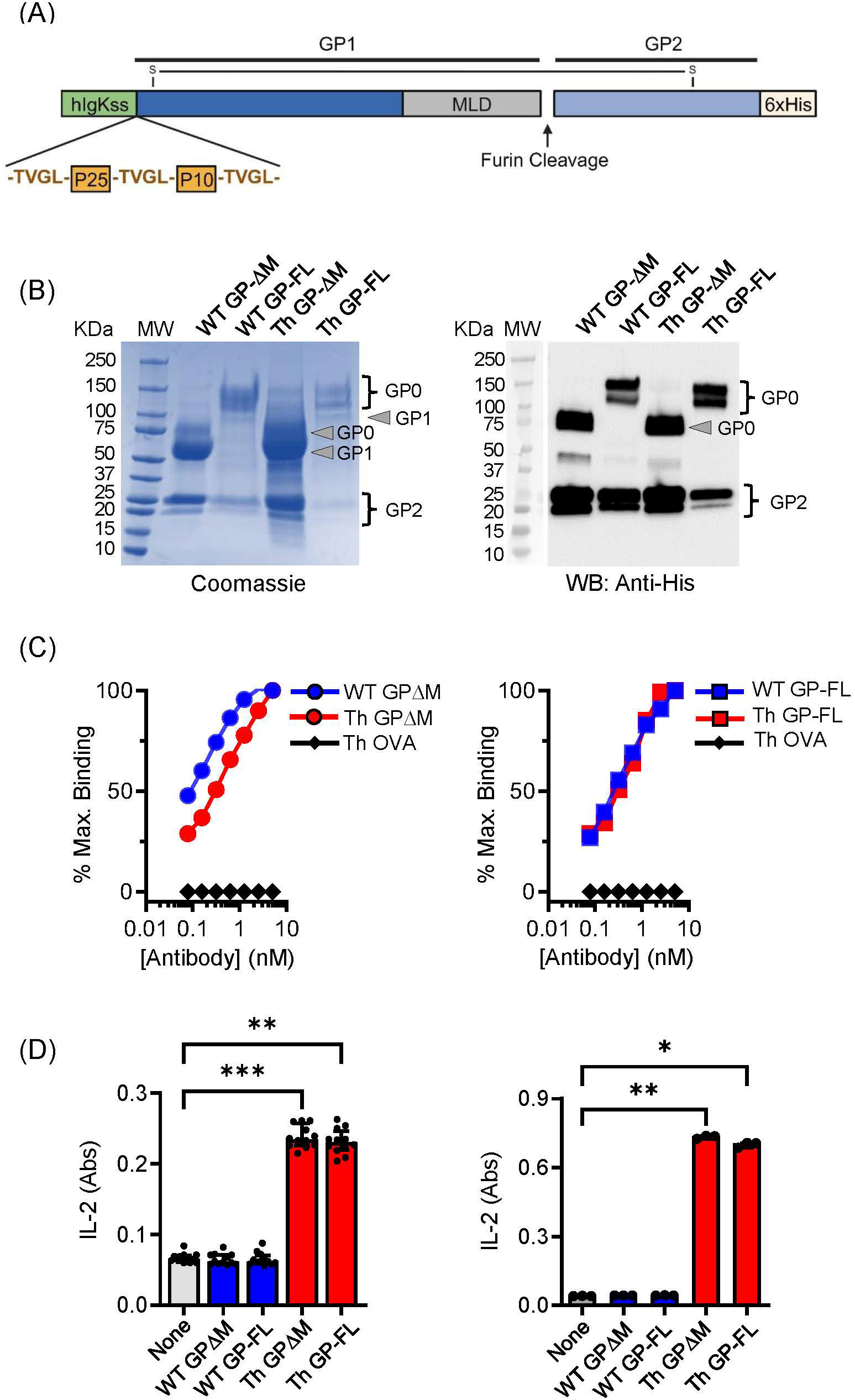
Characterization and comparison of EBOV GP vaccines. **(A)** Schematic of the Th GP-FL vaccine against EBOV. The Th vaccine consist of the N-terminal human Ig-kappa signal sequence (hIgKss), the BCG T helper epitopes (P25 and P10) which are flanked by the cathepsin B cleavage site (TVGL), GP1 which includes the mucin-like domain (MLD), the furin cleavage site, GP2, and the C-terminal hexahistadine tag (6xHis). GP1 and GP2 are held together by a disulfide bond. For the WT GP-FL version of the vaccine, the BCG T helper epitopes (P25 and P10) flanked by the cathepsin B cleavage sites are absent. The MLD deleted versions of these vaccines were also constructed (Th GPΔM and WT GPΔM). **(B)** SDS-PAGE under reducing conditions of purified EBOV GPs as shown by Coomassie gel staining and Western blotting with anti-His antibody. The GP1 and GP2 fragments, which are normally held together by disulfide bonds, were separately resolved under the reducing and denaturing conditions of the SDS-PAGE analysis. The expected size for GP2, which consists of the C terminal region of the EBOV GP after the MLD is 25 KDa. The expected size for the GP1 precursor is 120 KDa and 60 KDa for the full length and the MLD deleted version of the EBOV GP, respectively. The GP0 fragment in the full-length versions of the GP constructs was detected as two or more bands of ∼120-145 KDa, consistent with glycosylation and disordered structure of the MLD. Purity of isolated GPs was ≥ 90% based on Coomassie blue staining of the gels, and protein yields were determined by BCA protein assay (WT GP-ΔM: 2060 μg/ml, WT GP-FL: 220 μg/ml, Th GP-ΔM: 2972 μg/ml, Th GP-FL: 80 μg/ml). **(C)** ELISA with antibody ADI-15878 specific for EBOV GP conformational epitope was used to probe purified EBOV GPs. The ovalbumin version of the Th vaccine (Th OVA) served as a negative control to show the specificity of ADI-15878 antibody against EBOV GP. **(D)** Processing and presentation of BCG Th epitopes was shown by incubating purified EBOV GPs for 16 hours with dendritic cells and in the presence of a CD4^+^ T cell hybridomas specific for P25 of Ag85B (left) or P10 of TB9.8 (right). Supernatants were analyzed by sandwich ELISA for IL-2 (indicated as absorbance (Abs) values for conversion of the assay substrate. Multiple columns were analyzed by Kruskal-Wallis one-way ANOVA, followed by Dunn’s multiple comparison test; (***p < 0.001, **p < 0.01, *p < 0.05).

Protein purity and quality were assessed by SDS-PAGE and immunoblotting. As shown by Coomassie staining (Fig. 1B, left panel), the mature form (GP0) and the two proteolytic fragments (GP1 and GP2) of the full-length (WT GP-FL and Th GP-FL) and the ΔMLD versions (WT GP-ΔM and Th GP-ΔM) of EBOV GP constructs were observed and confirmed to be of the expected sizes ^40, 51^. As expected, immunoblotting with antibody specific for the hexahistidine tag at the carboxyl-terminal of the 25 kDa GP2 precursor (Fig. 1B, right panel) detected the mature form GP0 and the GP2 cleavage product, but not the GP1 fragment which lacks the histidine tag. Since EBOV GP exists mainly as trimers in its native cell surface form, we also carried out size exclusion chromatography to analyze monomeric versus multimeric state of the soluble GP constructs in solution ^52, 53^ (Supplemental Fig. 1). This showed retention times consistent with mass of 600 kDa or more for the proteins in solution, indicating complexes at least as large or larger than the expected size for soluble trimers. This suggested that the subunit vaccines produced here were likely to be a mixture of trimers and higher order multimers.

Consistent with the correctly folded structure for at least a fraction of the purified GP preparations, the conformation sensitive anti-EBOV GP antibodies ADI-15878 and KZ52 ^54^ bound to all of the purified proteins in solid phase ELISA, (Fig. 1C and Supplemental Fig. 2). To demonstrate correct processing and presentation of the BCG epitopes embedded in the Th (FL) and Th (ΔM) fusion proteins for T cell recognition, we used previously isolated mouse T cell hybridomas specific for the Ag85B or TB9.8 epitopes presented by MHC class II I-A^b^ molecules ^33, 55^. T cell hybridoma cells cultured with mouse bone marrow derived dendritic cells secreted IL-2 into the culture supernatants in response to the purified GPs containing the Th sequence encoding the relevant T cell epitopes, indicating efficient antigen processing at the inserted cathepsin S cleavage sites and presentation by I-A^b^ (Fig. 1D). Furthermore, the BCG epitopes incorporated into the Th vaccines were targeted by long-lived memory Th cells in BCG vaccinated mice. This was apparent in mice vaccinated with PBS or BCG and then rested for 17 weeks before administrating the Th GP-FL vaccine or PBS sham control. Two weeks later, IFNγ ELISPOT assays were performed on splenocytes to determine recall responses of BCG specific Th cell against the peptide-25 (P25) of the immunodominant Ag85B or Mtb (strain H37Rv) lysate (Figs. 2A and B). Mice in both of the BCG vaccinated groups developed BCG specific Th cells reactive to Mtb lysate, but only the BCG group that was subsequently immunized with the Th GP-FL vaccine showed significant expansion of P25 specific Th cells.

**Figure 2.**
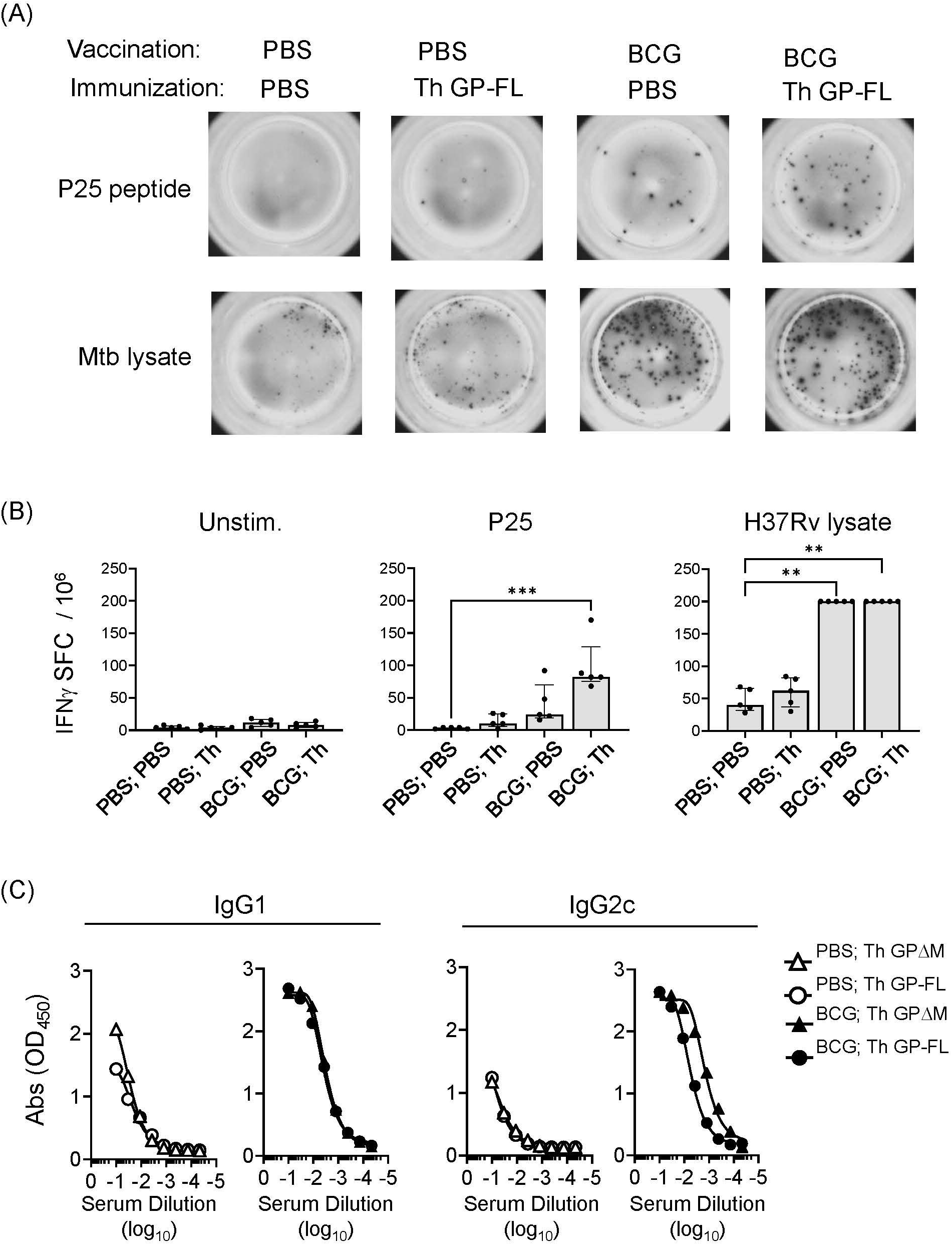
Induction of cellular and humoral immune responses by Th vaccines. For analysis of cellular responses, groups of mice (n = 5) were vaccinated with BCG or received sham vaccination with PBS injection, rested for 17 weeks and then immunized with the Th GP-FL or sham immunized (PBS). Two weeks after the immunization, IFNγ ELISPOT assays were performed on unstimulated, peptide-25 (P25) or Mtb (H37Rv lysate) stimulated splenocytes. (A) Representative spot forming cell (SFC) images of selected animals for each group. (B) Plots showing individual animal counts and group medians with interquartile range. Multiple columns were analyzed by Kruskal-Wallis one-way ANOVA, followed by Dunn’s multiple comparison test; (***p < 0.001 and **p < 0.01). Note that values for H37Rv lysate stimulation of BCG vaccinated groups are all plotted at the upper limit for accurate quantitation in the assay. (C) Mice (n = 5) were vaccinated with BCG or received sham vaccination with PBS injection, rested for 5 weeks and then prime and boosted with the EBOV GP vaccines (Th GP-FL or Th GPΔM). Two weeks after the boost, sera were collected, and antibody titers against EBOV GP (WT FL) were determined using ELISA specific for IgG1 or IgG2c isotypes.

To test the immunogenicity of the full length Th GP-FL vaccine and compare this directly with the ΔMLD version (Th GPΔM) that we previously showed to lower the vaccine dose required to induce antibody responses and induce IgG2c class switching ^33^, mice were vaccinated with BCG or received sham vaccination with PBS injection, and then primed and boosted by subcutaneous injections of either the Th GP-FL or Th GPΔM in alum. A solid phase ELISA was performed to detect the presence of anti-EBOV GP-IgG1 and -IgG2c antibodies. Confirming our previously published findings ^33^, the BCG-specific Th cells from prior BCG vaccination, which were absent in the sham vaccinated (PBS) groups, were recruited by the Th vaccine to promote class switching to IgG2c, and either version (Th GPΔM or Th GP-FL) of the Th vaccine induced similar antibody levels (Fig. 2C). Taken together, these results showed that the Th GP-FL can be used to replace the Th GPΔM version of the vaccine to more accurately represent the native form of GP associated with actual EBOV infection and provide the broadest array of potential epitopes for both neutralizing and non-neutralizing antibodies.

### BCG vaccination upregulates FcγRIV expression and supports long-lived antibody responses

Non-neutralizing antibodies mediate their functions primarily through the binding of FcγRs to recruit immune cell effector functions, including cytolysis and phagocytosis, to clear infected cells. In mice, the Fc portions of the IgG2 isotypes have the highest affinities for FcγRIV, which is abundantly expressed on monocytes, macrophages, and neutrophils ^31^, and to a lesser degree on NK cells ^56, 57^. To determine the expression level of FcγRIV on immune cells, flow cytometry was performed to identify B cells (B220^+^) NK cells (NK1.1^+^), neutrophils (CD11b^high^ Ly6G^high^), macrophages (CD11b^high^ Ly6C^high^) and monocytes (CD11b^high^ Ly6C^low^) (Fig. 3A). At 2 weeks after BCG vaccination, an increase in the levels of FcγRIV was detected on monocytes, macrophages, NK cells, and B cells (Fig. 3B). Although the Th vaccine expanded the BCG memory Th cells (Fig. 2), this was not associated with further enhancement of the BCG induced FcγRIV expression at 17 weeks after the initial BCG vaccination (Fig. 4A & B). However, these findings showed that BCG vaccination induced prolonged elevation of FcγRIV expression on effectors cells, which is likely to be relevant to the efficacy of the Th vaccine design that favors the induction of IgG2c class-switched antibodies ^33^.

**Figure 3.**
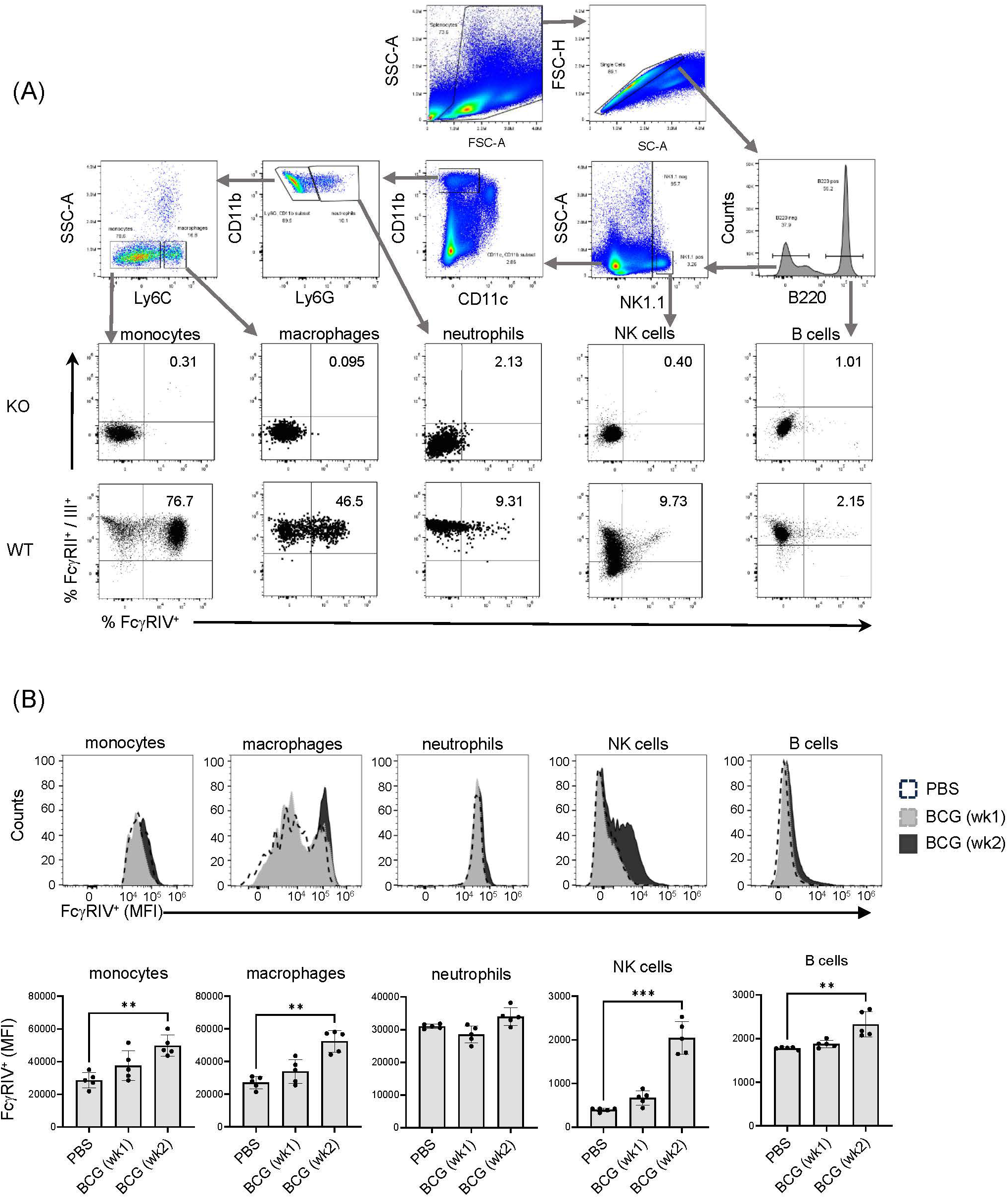
Expression of FcγRIV increases at week 2 after BCG vaccination. **(A)** Flow cytometry gating strategy using mice with compound genetic knock out of FcγRII, RIII and RIV α-chains (KO) and wildtype (WT) C57BL/6 mice to determine the gating for FcγRII/III and FcγRIV expression on immune cells. After singlet cell gating, the corresponding surface markers were used to stain splenocytes to identify the following immune cells: monocytes (CD11b^+^ Ly6C^low^), macrophages (CD11b^+^ Ly6C^high^), neutrophils (CD11b^+^ Ly6G^+^), NK cells (NK1.1^+^), and B cells (B220^+^). **(B)** Mice (C57BL/6) were vaccinated with 10^7^ BCG per mouse or received PBS injections as control. Spleens were harvested at week 1 after BCG vaccination (gray histogram), or at week 2 after PBS injection (white histogram) or BCG (black histogram) vaccination, and splenocytes were analyzed by FACS to determine the expression level of FcγRIV. Top panel shows representative histograms for an individual mouse from each group, and bottom panel shows median of MFI values for 5 mice in each group on each indicated cell type. Median with interquartile range for five replicates is shown and results were analyzed using Kruskal-Wallis one way ANOVA nonparametric test and Dunn’s multiple comparison test; (***p < 0.001, **p < 0.01).

**Figure 4.**
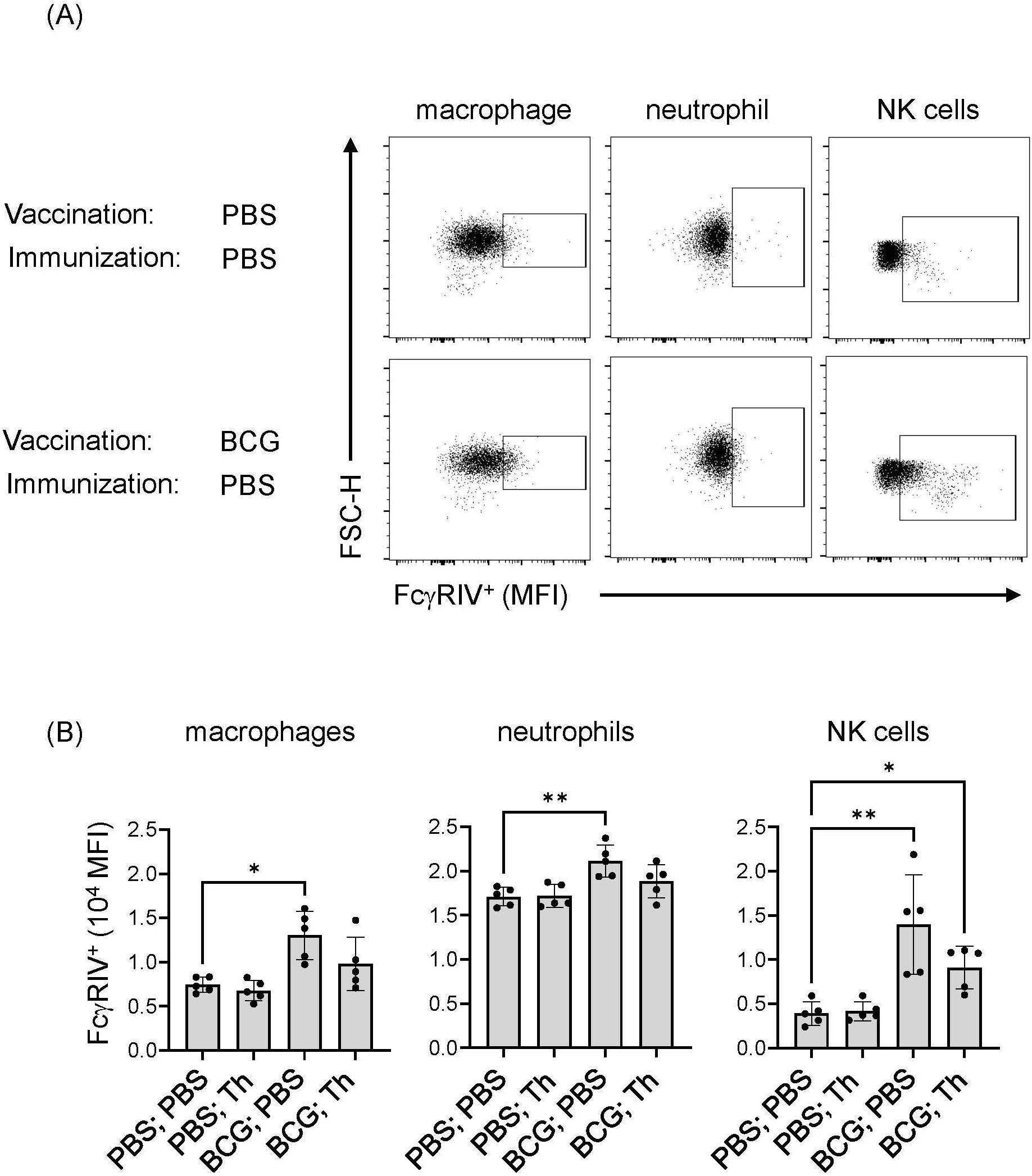
BCG-induced FcγRIV expression was maintained after Th vaccination. Mice (n = 10) were sham vaccinated with PBS or with BCG at 10^7^ CFU per mouse and rested for 17 weeks. Each group was further subdivided into 2 groups (n =5) and received either PBS or the Th GP-FL vaccine at 0.05 μg per mouse in alum, and splenocytes were harvested 2 weeks later to determine the level of FcγRIV expression by FACS. Singlet gating on splenocytes were stained for macrophages (CD11b^high^ Ly6C^high^), neutrophils (CD11b^high^ Ly6G^high^), and NK cells (NK1.1^high^) for expression of FcγRIV (CD16.2) as shown in **(A)** for representative animals of sham vaccinated with PBS and with BCG alone, and quantified in **(B)** by MFI levels for the 5 animals in each group. The median with interquartile is shown and analyzed using Kruskal-Wallis one way ANOVA nonparametric test with and Dunn’s multiple comparison test; (**p < 0.01 and *p < 0.05).

To determine the duration of the persistence of antibodies against EBOV GP in mice receiving the Th GP-FL vaccine, mice were either vaccinated with BCG or sham vaccinated (PBS only), and then immunized with 5 μg of Th GP-FL (Fig. 5). In this experiment, a higher dose of the Th GP-FL was given to the animal instead of the usual dose of 0.5 μg per mouse in order to elicit a detectable IgG2c response in the PBS group for comparison with the BCG group. Serum samples were collected at times ranging from 2 to 39 weeks after the administration of the Th GP-FL vaccine and analyzed by ELISA for anti-EBOV GP titers for both IgG1 and IgG2c subclasses. Compared to the PBS group that lacked BCG specific Th1 cells, the intrastructural help provided by BCG specific Th1 cells in the BCG vaccinated group promoted higher anti-EBOV GP titers. Anti-EBOV GP antibodies were detected even at week 39 after vaccination (Fig. 5A), and, at the same time, plasma cells secreting these anti-EBOV GP antibodies were detected in bone marrow and not the spleen (Fig. 5B), indicating that long lived plasma cells induced by the Th vaccine can elicit long lasting protection.

**Figure 5.**
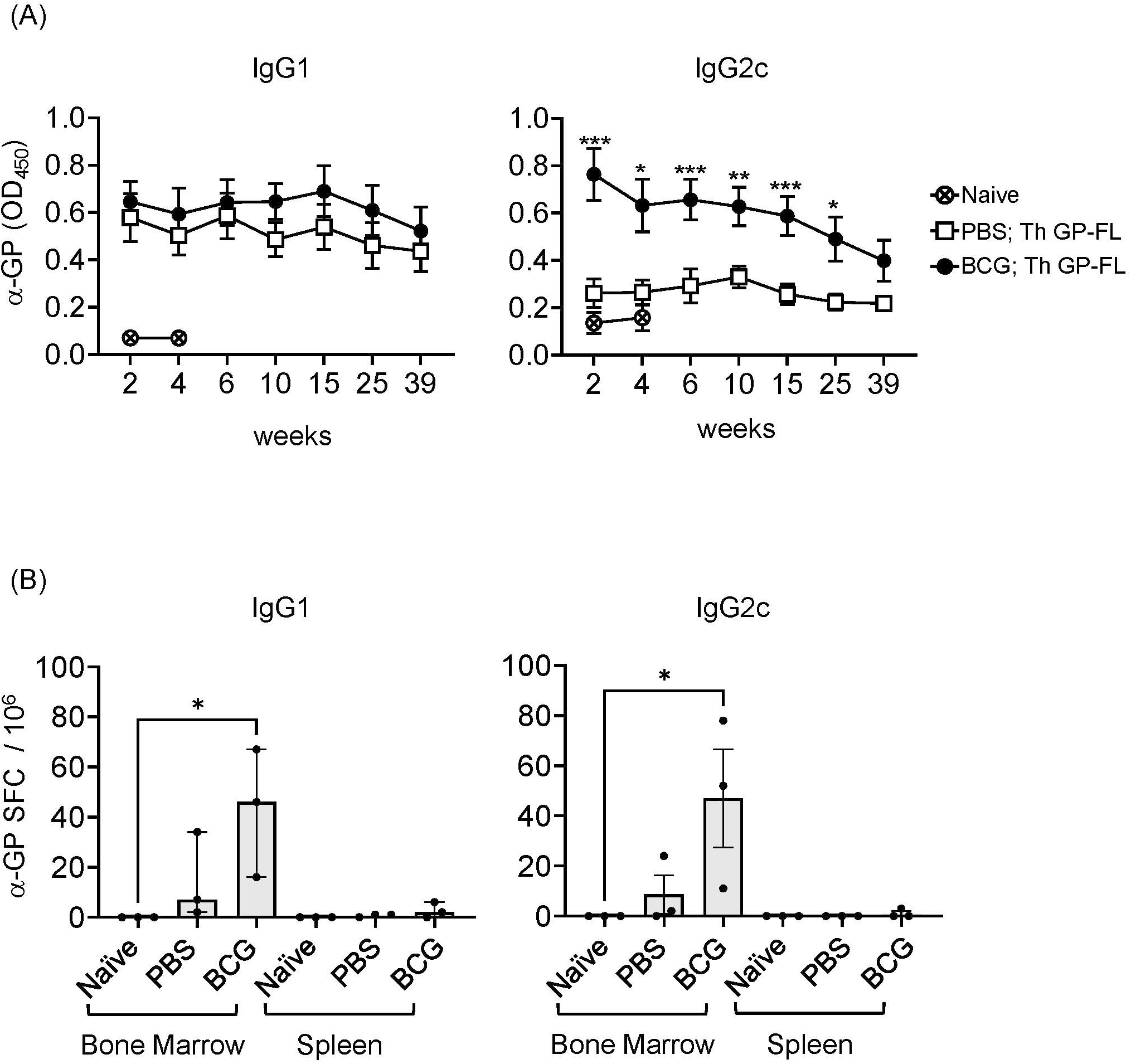
Long lasting humoral immune response induced by the Th vaccine. **(A)** Serum samples from sham PBS (□) or BCG (•) vaccinated mice and subsequently immunized once with 5 μg of the Th GP-FL vaccine were analyzed by ELISA for anti-EBOV GP IgG1 (left panel) or IgG2c (right panel) antibodies throughout the time course of 39 weeks. Serum from a naïve mouse (⊗) collected at week 2 and 4 was used to measure the background level for the ELISA assay. **(B)** Splenocytes and bone marrow cell suspensions from these mice at 39 weeks were harvested and used for B cell ELISPOT assay to quantitate anti-EBOV GP IgG1 or IgG2c antibody secreting spot forming cells (SFCs). The median values with interquartile ranges are shown and analyzed using Kruskal-Wallis one way ANOVA nonparametric test with Dunn’s multiple comparison test; (*p < 0.05).

### BCG vaccination induced extrafollicular Th1 responses and altered germinal center formation

In our previous publication, we showed that antibodies induced by the Th vaccine have different affinities in the Fab region that correlated with IgG1 and IgG2c isotypes ^33^. B cells that enter the germinal center (GC) form cognate interaction with T follicular helper (Tfh) cells, which are defined by expression of CXCR5 and the lineage-defining transcription factor Bcl-6 ^58^, and go through multiple rounds of affinity maturation to develop high affinity antibodies. B cells that encounter antigens outside of GCs undergo cognate interactions with non-Tfh cells such as Th1 cells, which reduces affinity maturation but provides rapid protection in early stages of infection ^59^. To visualize the location of BCG-specific Th cells within a secondary lymphoid organ after BCG vaccination, we used adoptive transfer of GFP labelled CD4^+^ T cells expressing a TCR transgene specific for the P25 epitope of BCG Ag85B as previously described ^33^. To compare the extent to which these adoptively transferred T cells remained extrafollicular or were capable of entering germinal centers, we compared mice vaccinated with BCG versus mice immunized with P25 peptide combined with various adjuvants, including alum and a multicomponent formulation known as LASTS-C (lipid A, Span8, Tween 80 and CpG oligodeoxynucleotides) ^43^. 16 hours after immunization, splenocytes were isolated and analyzed by FACS with gating on CD4^+^ GFP^+^ cells (Fig. 6A; top panel). Th1 polarization of the transferred P25-specific GFP^+^ CD4^+^ T cells as determined by Tbet expression was strongest in BCG vaccination as compared with other adjuvants (Fig. 6A; middle panel), whereas the Tfh polarization as shown by CXCR5 and Bcl-6 double staining was extremely low except in animals receiving vaccination with the LASTS-C adjuvant (Fig. 6A; lower panel), which correlates with its ability to induce strong neutralizing antibody responses ^44^.

**Figure 6.**
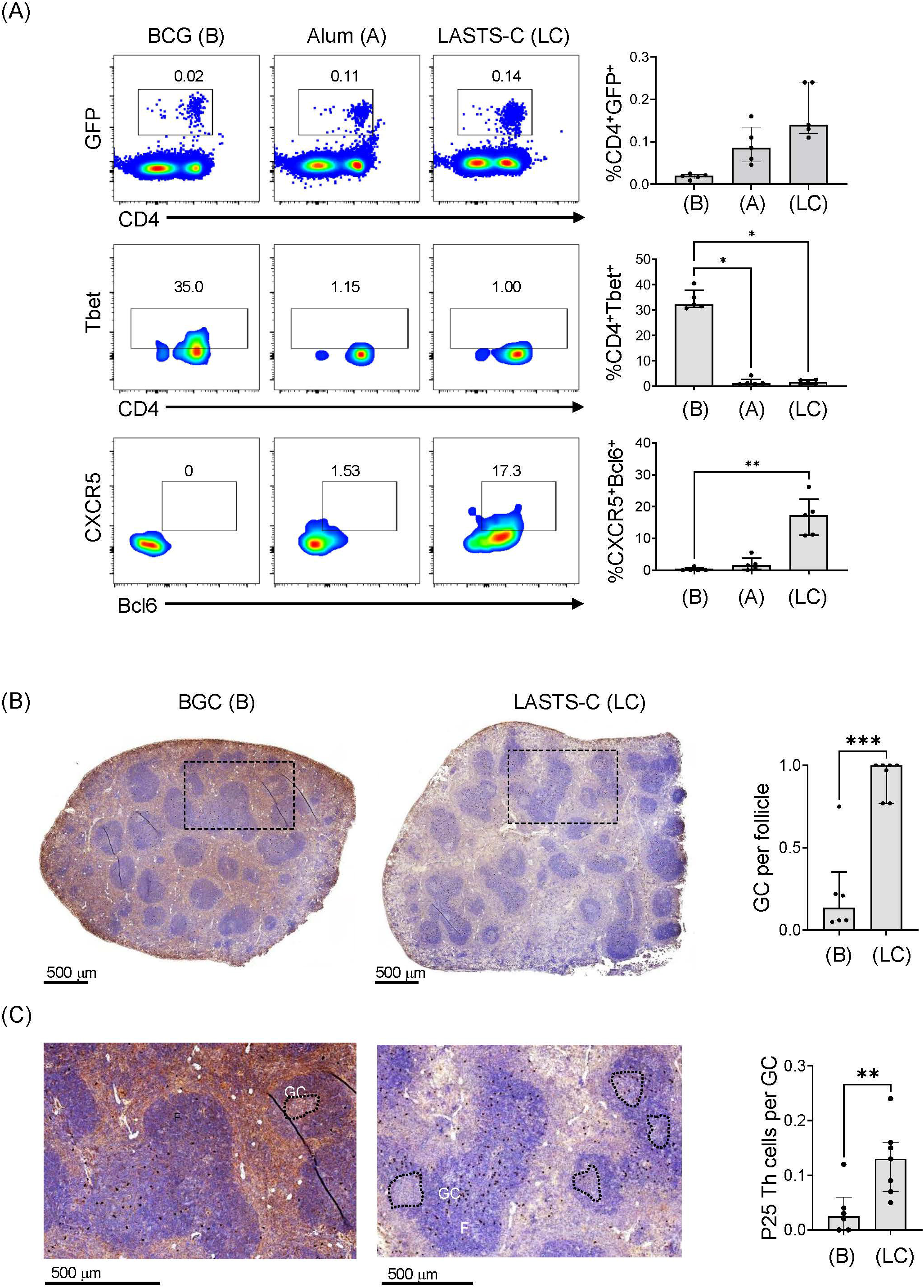
BCG vaccination promotes predominantly Th1 responses. **(A)** Wildtype mice were adoptively transferred with 4 x 10^4^ T cells purified from P25 TCR-Tg GFP^+^ mice 16 hours prior to vaccination with BCG (B), or peptide-25 adjuvanted with either alum (A) or LASTS-C (LC). The mice were sacrificed 6 days after vaccination, and splenocytes (n=5) were analyzed by FACS for master transcription regulators and key markers for Th1 (Tbet) and Tfh (CXCR-5 and Bcl-6). Multiple comparisons were analyzed by Kruskal-Wallis one-way ANOVA (**p < 0.01, ***p < 0.001). **(B)** Wildtype mice were adoptively transferred with 4 x 10^4^ T cells purified from P25 TCR-Tg GFP^+^ mice 16 hours prior to vaccination with BCG or with the P25 peptide adjuvanted with LASTS-C. Formalin-fixed and paraffin-embedded spleens were cut into thin sections for immunocytochemistry with anti-GFP followed by hematoxylin and eosin counterstaining. **(C)** High magnification of boxed areas in (B) to visualize the P25 specific T cells as dark colored spots. Follicles (F), germinal centers (GC), and P25 Th cells were quantified manually by a blinded observer as the number of GC per follicle in (B), and the number of P25 Th cells per GC in (C). Medians with interquartile ranges for 6-7 sections derived from 3 mice from each group are plotted. Mann-Whitney test was used for pairwise comparison (**p < 0.01, ***p < 0.001).

To further evaluate the effects of BCG vaccination on the functional outcomes of CD4^+^ T cell responses, we analyzed the localization of P25 specific T cells in the spleen by immunohistochemistry. Naïve P25-specific GFP^+^ CD4^+^ T cells were transferred intravenously into mice that were vaccinated 16 hours later with either BCG or the P25 peptide adjuvanted in LASTS-C. Six days after vaccination, spleens were isolated, sectioned, and analyzed by immunohistochemistry with anti-GFP staining followed by H&E counter staining (Fig. 6B). BCG vaccination, as expected for strong Th1 biasing stimuli, diminished the formation of GCs (Fig. 6B; left panel) as compared to non-Th1 adjuvant such as LASTS-C (Fig. 6B; right panel), which was quantified by counting the number of GC per follicle (Fig. 6B). In BCG vaccinated mice, the GFP^+^ CD4^+^ T cells, observed as brown precipitate of the 3,3-diaminobenzidine in the immunohistochemistry staining with anti-GFP conjugated with horse radish peroxidase, were present in the white pulp areas (Fig. 6B; left panel) but nearly absent in the relatively scarce GCs (Fig. 6C; left enlarged panel). In contrast, in the LASTS-C vaccination group (Fig. 6B; right panel) there were numerous GCs, and the GFP^+^ CD4^+^ T cells were mostly localized within the GCs (Fig. 6C; right enlarged panel), as quantified as the number of P25 cells per GC (Fig. 6C). These data suggested that the majority of BCG Th cells remained outside the GC at the border of the B cell follicle and likely exerted their effects on the antibody response at this location comprising the boundary of B and T cell zones.

### Vaccination with subunit vaccine for protection against EBOV challenge

Experiments to assess our proposed regimen for protection from lethal EBOV infection required the use of mice that will have reached 24 weeks of age at the time of infection with EBOV, an age group that has not to our knowledge been previously tested in the EBOV challenge model. To determine whether mice at this age have similar susceptibility to EBOV infection compared to the typically used younger mice, 24-week-old mice were challenged with 10 or 1,000 focus-forming units (FFU) of mouse-adapted EBOV (MA-EBOV). Eight days after challenge, 24-week-old mice succumbed to the MA-EBOV infection even with the lower 10 FFU infectious dose (Supplemental Fig. 3), which was similar to the time to death observed previously in younger mice at 6-14 weeks of age ^46^. Having established the susceptibility of the older mice in this model, we tested the efficacy of our regimen using recombinant protein vaccines augmented by prior BCG vaccination to protect against EBOV challenge using the vaccination strategy illustrated in Figure 7A. Sera were analyzed by ELISA for anti-EBOV GP antibody responses 2 weeks after priming and again after boosting with the subunit vaccines. The group receiving the WT GP-FL vaccine (WT), which is incapable of recruiting BCG Th cells for intrastructural help, failed to induce robust anti-EBOV GP antibody responses, and did not undergo IgG2c class-switching (Fig. 7B, left). This was in contrast to the Th GP-FL vaccine (Th), which enhanced anti-EBOV GP antibody responses and IgG2c class-switching (Fig. 7B, right). An ELISA performed with end-point dilution of the serum samples collected after the Th-vaccine boost showed similar results, with the Th GP-FL vaccine providing elevated levels of both IgG1 and IgG2c antibodies against EBOV GP (Fig. 7C). Five weeks after boosting with the subunit vaccines, mice were challenged with 10 FFU of MA-EBOV. All the mice from the PBS and the WT GP-FL vaccine group succumbed to the infection 8 days after MA-EBOV challenge, whereas the majority of the mice in the Th GP-FL vaccine group survived through the end of the study, at which point they appeared healthy and had regained their original body weight (Fig. 7D). In addition, at day 5 after EBOV infection, five mice were sacrificed to determine the viral titers in the blood and organs. Mice that received the Th GP-FL vaccine had a lower viral titer in the blood, liver, and spleen when compared to mice that either received no vaccination or the WT GP-FL (Fig. 7E). An ELISA was also performed on the serum samples collected from these mice which showed that mice that received the Th GP-FL vaccine also had a higher anti-EBOV GP antibody titer compared to the control group receiving PBS only or the WT GP-FL vaccine group (Fig. 7F). Long term survivors from the Th vaccine group were also bleed at termination of the experiment (day 112, corresponding to 14 days after EBOV challenge), and analysis of these serum samples showed persistently high levels of anti-EBOV GP antibodies (Fig. 7G). Thus, the Th vaccine strategy clearly protected the mice against lethal EBOV infection by limiting viral replication to control the early stage of infection, which is known to be important in conferring protection as seen in other viral infections ^59^.

**Figure 7.**
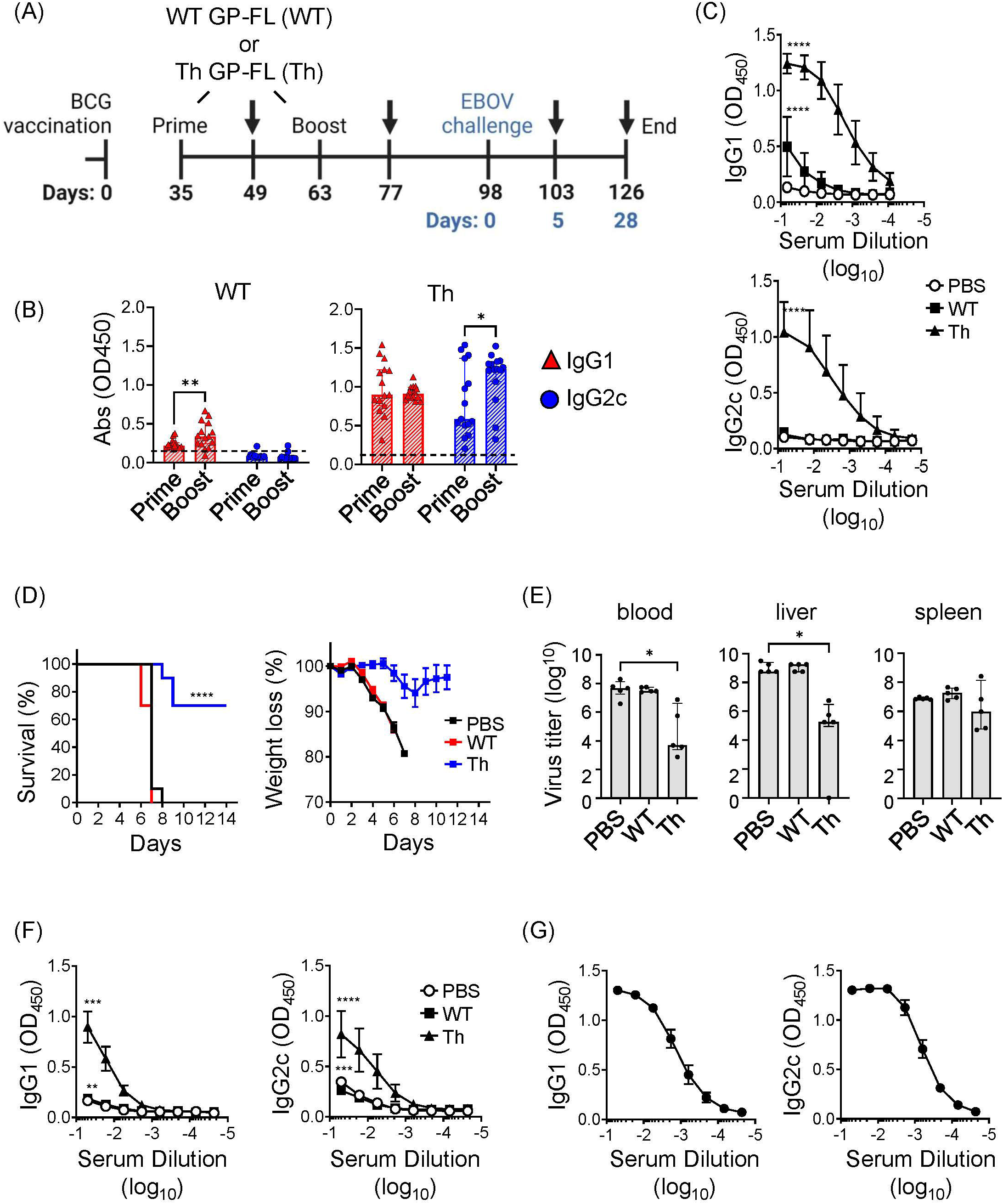
Vaccine induced protection against lethal EBOV infection. **(A)** Timeline of the EBOV challenge experiment. C57BL/6NHsd mice (n = 15) were vaccinated with BCG, and subsequently primed and boosted with 0.05 μg of the indicated recombinant protein vaccine (WT GP-FL or the Th GP-FL) in alum as indicated in the schematic. MA-EBOV infection (i.p.) with 100 pfu per mouse was performed on day 98, which corresponds to day 0 (blue) of the challenge experiment. Arrows represent blood sample collections, and harvesting of blood, livers, and spleens at the last collection point after sacrifice. **(B)** Serum samples collected 2 weeks after administering WT GP-FL (WT) or the Th GP-FL (Th) vaccines for both priming and boosting time-points were analyzed by ELISA for anti-EBOV GP IgG1 (red) and IgG2c (blue) antibodies. **(C)** Endpoint titers of anti-EBOV GP antibodies following boosting with the WT or Th vaccines were also analyzed by ELISA in sera from mice primed and boosted with either PBS or 0.05 μg of the indicated recombinant protein vaccine. **(D)** Naïve mice (black) or mice that received the WT (red) or Th (blue) vaccines were challenged with MA-EBOV and the times to death (left panel) and body weight changes (right panel) for individual mice were recorded. The Mantel-Cox test was performed for comparison of the survival curves; (****p < 0.0001). **(E)** Five days after the challenge, mice (n = 5) were sacrificed to determine viral titers from the corresponding organs. The median with interquartile ranges is shown and analyzed using Kruskal-Wallis one way ANOVA nonparametric test with and Dunn’s multiple comparison test; (**p < 0.01, *p < 0.05). (**F**) The anti-EBOV GP IgG1 and IgG2c antibody titers were also measured from serum of these mice sacrificed at day 5 after challenge (****p < 0.0001, **p < 0.01, *p < 0.05 two-way ANOVA test and Sidak’s multiple comparison test). (**G**) Twenty-eight days after the challenge, the surviving mice were sacrificed, and blood was collected to measure anti-EBOV GP IgG1 and IgG2c antibody titers.

The approach to antiviral vaccination used in the current study is based on the classic hapten-carrier immunization studies that led to the understanding of the concept of linked recognition. Many effective vaccines depend on the core immunological concept of linked-recognition, in which Th cells recognize processed peptides derived from the immunogen targeting B cell receptors to provide intrastructural help to B cells, leading to T cell dependent antibody responses ^60^. These include vaccines that rely on the production of antibodies against targets that entirely lack T cell epitopes, such as those against *Haemophilus influenzae* type b (Hib) polysaccharides ^61^ or small hapten-like molecules like nicotine ^62^. In these cases, conjugation to a protein carrier containing Th cell epitopes is required to elicit an optimal T cell dependent B cell response. In a logical extension of this principle, we and others have applied this approach to creating protein subunit vaccines against viral antigens to enhance and accelerate protective antibody responses through the recruitment of pre-existing Th cells against other potent antigens, such as those delivered by previous vaccination against pathogens such as mycobacteria. For example, Klessing et al. developed a vaccine against HIV that can recruit intrastructural help from Th cells induced by an *M. tuberculosis* subunit vaccine, and showed that this approach induced higher antibody titers that persisted for extended period of time ^63^. In our previous work we applied a similar approach to capture intrastructural help to B cells from pre-existing Th1 cells specific for immunodominant mycobacterial antigens in BCG vaccinated mice ^33^. In the current study, we expanded on our previous work to determine the protective efficacy of this Th vaccine design against EBOV challenge in the mouse model, and to explore in greater detail the potential mechanisms mediating this protection. Consistent with our findings, we showed that this vaccine strategy induced antibody class-switching to IgG2c, an isotype that is known to have high affinity toward FcγRIV, suggesting that antibodies with effector activities such as antibody-mediated cellular cytotoxicity (ADCC) might be a key feature that extended the antiviral effects beyond simple neutralization of viral entry.

Our previous efforts to generate fusion proteins for use as subunit vaccines against EBOV used a truncated form of EBOV GP in which the MLD was deleted, which our preliminary work had shown to be produced with much higher yields than a full-length version (Th GP-FL) that retains the MLD ^33^. In the current study, we further improved production and purification of the Th GP-FL fusion protein and formulated this with alum to generate a candidate vaccine against EBOV infection and disease for use in previously BCG vaccinated hosts. This full-length version of the extracellular domains of EBOV GP should present an immunogen that corresponds more closely than the previously designed Th GPΔM vaccine that lacks the MLD to the actual infectious virus or the form of the EBOV GP expressed on the surface of infected host cells, making it potentially more effective for generating a broad range of antibodies mediating a variety of host protective functions. In this regard, while epitopes in the MLD region have not been strongly associated with broad neutralization of viral entry, such antibodies may play an important role in controlling the progression and spread of infection through non-neutralizing activities such as ADCC ^64^.

Our preparations of the Th GP-FL fusion protein produced so far appeared to exist mainly as higher order multimers in solution, rather than as soluble monomers or native homotrimers (Supplemental Fig. 1). The multimerization of the protein may be due to artifactual disulfide bond formation or other tight interactions that formed during the purification, and suggests the need for further optimization of the production and purification process. However, irrespective of the presence of larger multimeric complexes in the Th GP-FL preparations, the native conformation of the protein appeared to be present at significant levels, as shown by its recognition by the anti-EBOV GP monoclonal antibodies, ADI-15878 and KZ52, which recognize conformational epitopes of the native protein (Fig. 1C and Supplemental Fig. 2). Furthermore, the Th GP-FL vaccine was able to induce anti-EBOV GP antibodies that recognized EBOV GP expressed on the surface of recombinant vesicular stomatitis virus (Figs. 2C, 5A, and 7B). Most importantly, the vaccine conferred protection against challenge with MA-EBOV (Fig. 7D & E), indicating that antibodies generated by the Th GP-FL vaccine, particularly when administered in the context of prior BCG vaccination, were able to recognize the relevant form of GP during viral infection.

A key feature of our vaccine strategy is the prior BCG vaccination, which not only induced memory BCG-specific Th1 cells to provide intrastructural help, but also through trained immunity, can enhance non-specific immune mechanisms ^65^. Protection from trained immunity induced by BCG has been described in COVID-19 infection and also in BCG-based bladder cancer treatments ^66^. In our analyses, we observed that BCG exposure also promoted FcγRIV expression (Figs. 3 and 4) and IgG2c class-switching (Fig. 2C), which can be viewed as additional aspects of trained immunity. In the mouse model, the IgG2c isotype and FcγRIV expression together are important for the induction of ADCC by certain immune effector cells such as neutrophils and NK cells. Correlating with the induction of anti-EBOV GP IgG2c antibodies together with FcγRIV expression, BCG-vaccinated mice that received the Th GP-FL vaccine, but not those with the WT GP-FL vaccine, survived the EBOV challenge (Fig. 7D). This suggests that the anti-EBOV GP IgG2c isotype (Fig. 7B) played a significant role in conferring this protection. Analyzing the location of the BCG Th1 cells revealed that the anti-EBOV GP IgG2c antibodies were likely derived from extrafollicular plasmablasts since BCG vaccination induces a strong Th1 cell response that favor less GC development as compared to a Tfh promoting adjuvant (Fig. 6). As a result of this massive Th1 polarization, most of the T-dependent B cells are activated by the Th1 cells at the boundary of B cell follicles and not inside GCs. These GC-nonresident B cells develop into extrafollicular plasmablasts which are usually short-lived. Surprisingly, at 39-weeks after vaccination anti-EBOV GP antibody levels were still detected in vaccinated mice (Fig. 5A) and the EBOV GP-specific antibody secreting cells were still detected in the bone marrow (Fig. 5B), presumably representing long-lived plasma cells. This suggests the possibility that GC in BCG vaccinated mice, although not initially detected, may form at a later time and enable extrafollicular B cells induced in early stages post-vaccination to develop into memory B cells and long-lived plasma cells that take up residence in the bone marrow to sustain long term production of anti-EBOV GP antibodies.

Although other forms of EBOV vaccine are available such as the VSV-based EBOV vaccine (Ervebro) that is FDA approved for human use ^67–69^ and confers protection in non-human primates 10 days after vaccination ^70^, the disadvantages faced by virus vectored EBOV vaccines are manufacturing difficulties for large scale production and cold chain requirement during distribution ^69^, which can easily overwhelm logistics when dealing with larger outbreaks. Other EBOV vaccine platforms require strong adjuvants to increase immunogenicity ^49, 50, 71^, including recombinant subunit vaccines based on EBOV GP that are currently being developed ^49, 50, 71^. The development of a subunit EBOV vaccine should allow easier production and distribution, especially in resource limited nations, which can contribute to rapid deployment to control outbreaks ^72^. The recombinant protein vaccine used in this study has the unique ability to recruit BCG-specific Th1 cells to provide intrastructural help for driving antibody production against the recombinant protein subunit without the use a strong adjuvant that can increase cost and unwanted side effects. These properties can also lower the dose of the recombinant vaccine required, which can have an impact on manufacturing, cost, and distribution worldwide. Furthermore, sustained antibody responses through week 39 was observed after administering a single dose of the Th GP-FL vaccine. The benefit of the Th GP-FL vaccine developed in this study as a recombinant protein, no doubt, is its simplicity as compared to a virus vaccine, and its ability to harness pre-existing BCG-induced immunity to protect mouse against MA-EBOV infection. Based on current projections ^73^, BCG vaccination will continue in many regions of the world well into the future ^74^, thus establishing large populations that should be well suited for mass vaccination against EBOV or other emerging viruses ^75^ using the approach demonstrated by the current study.

## 5 Conflict of Interest

The authors declare that the research was conducted in the absence of any commercial or financial relationships that could be construed as a potential conflict of interest.

## 6 Author Contributions

TWN and SAP: experimental conception, design, analysis, interpretation of data, and writing of the manuscript. TWN: performed experiments and the analysis and acquisition of data. WF and AM: design, performed, acquired, and analyzed data for the EBOV mouse challenge. ASW: prepared the rVSV EBOV GP. NASA: assisted with analysis of histology data of spleen sections. CTJ: assisted with the FACS analyses. AM, WRJ, and KC: analyzed, interpreted experiments, and reviewed the manuscript. All authors critically reviewed and approved the manuscript.

## 7 Funding

The core facilities used in this study were all supported in part by NCI Cancer Center Service Grant P30CA013330. Shared instrumentation grants funded the purchase of the Cytek Aurora FACS analyzer (S10OD026833-01) and the 3DHistec Panoramic 250 Flash II slide scanner (1S10OD019961-01) used in this study. This study was in part supported by the Intramural Research Program, NIAID, NIH (AM).

## 8 Acknowledgments

Resources and advice were provided by core facilities at Albert Einstein College of Medicine, including the Flow Cytometry, Analytic Imaging and Histopathology facilities. We thank Dr, Scott Garforth and the Macromolecular Therapeutics Development Facility at Albert Einstein College of Medicine for performing the size exclusion chromatography. We also thank Mei Chen and John Kim (Department of Microbiology & Immunology, Albert Einstein College of Medicine) for expert technical assistance with mouse experiments. We thank Bing Chen (Department of Microbiology & Immunology, Albert Einstein College of Medicine) for assistance in maintaining of FcγR KO mice colony. We also thank the animal care takers of the Rocky Mountain Veterinary Branch (NIAID, NIH) for their support of the EBOV mouse challenge.

## 11 Supplementary Material Captions

**Supplemental Figure 1. Analysis of EBOV GP by size exclusion chromatography.** Purified wildtype EBOV GP full-length (WT GP-FL) and the Th EBOV GP vaccine (Th GP-FL) were subjected to the SRT SEC-300 size exclusion column. SRT SEC-300 exclusion limit is 1250 kDa.

**Supplementary Figure 2. Comparison of EBOV GP vaccines. (A)** ELISA with antibody KZ52 specific for EBOV GP conformational epitopes was used to probe purified EBOV GP. The ovalbumin version of the Th vaccine (Th OVA) served as a negative control to show the specificity of KZ52 antibody against EBOV GP (left panel).

**Supplemental Figure 3. Susceptibility of 24-week old C57BL/6N to MA-EBOV challenge.** 24-week old mice were infected with 10 or 1000 FFU of mouse-adapted (MA-) EBOV and succumbed to infection by day 7-8 after EBOV challenge.

## Notes

### Competing Interest Statement

The authors have declared no competing interest.

