## Supplementary figures and images for "A viral vaccine design harnessing prior BCG immunization confers protection against Ebola virus"

### Supplemental Figure 1

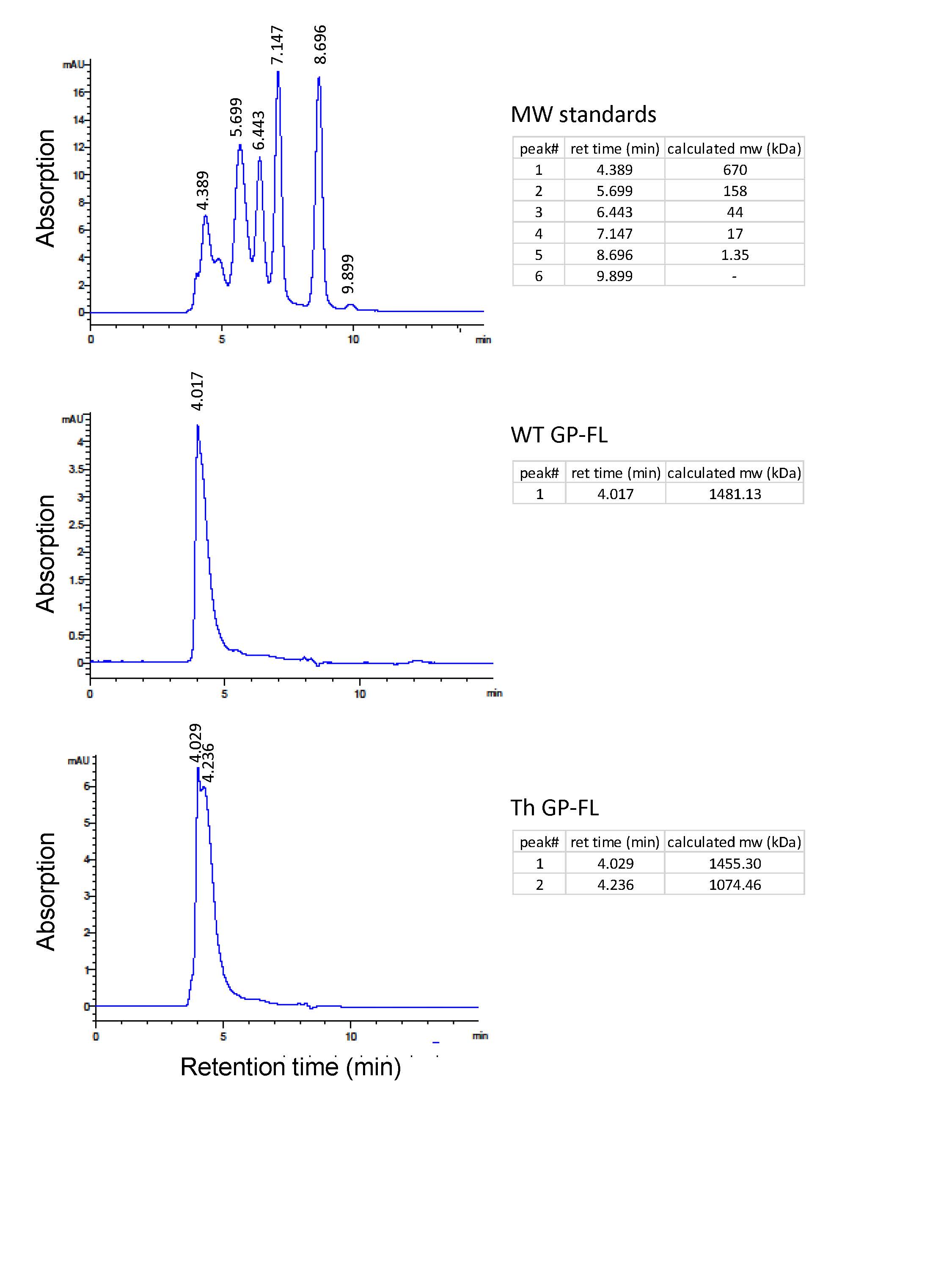

### Supplemental Figure 2

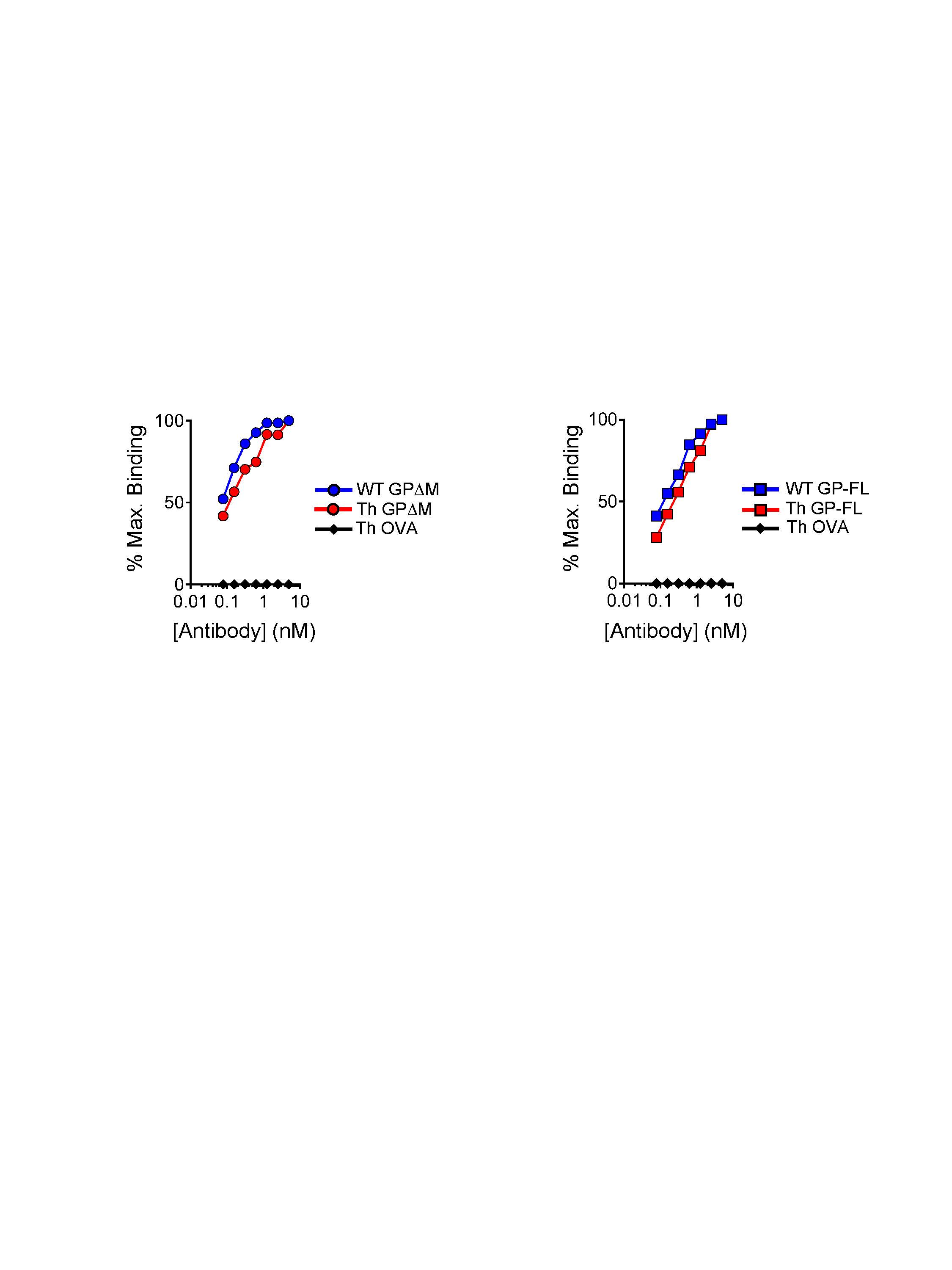

### Supplemental Figure 3

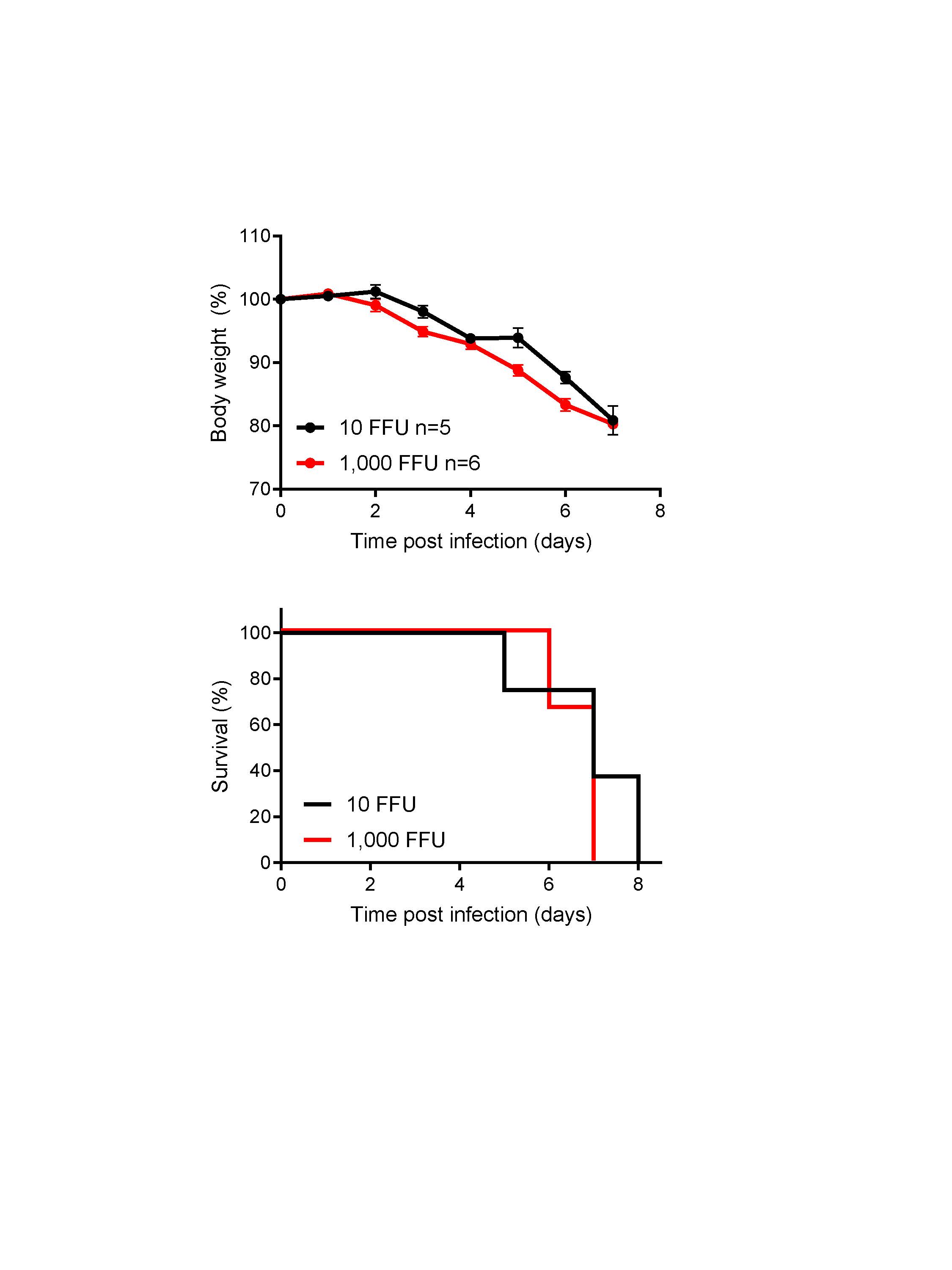
